# Pvr and downstream signaling factors are required for spreading of *Drosophila* hemocytes at larval wound sites

**DOI:** 10.1101/2021.06.10.447972

**Authors:** Chang-Ru Tsai, Alec Jacobson, Niki Sankoorikkal, Josue D. Chirinos, Sirisha Burra, Yan Wang, Nishanth Makthal, Muthiah Kumaraswami, Michael J. Galko

**Affiliations:** Program in Developmental Biology, Baylor College of Medicine, Houston, Texas, United States; Department of Genetics, University of Texas MD Anderson Cancer Center, Houston, Texas, United States; Department of Pathology and Genomic Medicine, Houston Methodist Hospital, Houston, Texas, United States; Genetics & Epigenetics Graduate Program, University of Texas MD Anderson Cancer Center, Houston, Texas, United States

**Keywords:** inflammation, larvae, cell-spreading, Pvr, hemocytes, *Drosophila*, wound closure

## Abstract

Tissue injury is typically accompanied by inflammation. In *Drosophila melanogaster*, wound-induced inflammation involves adhesive capture of hemocytes at the wound surface followed by hemocyte spreading to assume a flat, lamellar morphology. The factors that mediate this cell spreading at the wound site are not known. Here, we discover a role for the Platelet-derived growth factor (PDGF)/ Vascular endothelial growth factor (VEGF)-related receptor (Pvr) and its ligand, Pvf1, in blood cell spreading at the wound site. Pvr and Pvf1 are required for *spreading in vivo* and in an *in vitro* spreading assay where spreading can be directly induced by Pvf1 application or by constitutive Pvr activation. In an effort to identify factors that act downstream of Pvr, we performed a genetic screen in which select candidates were tested to determine if they could suppress the lethality of Pvr overexpression in the larval epidermis. Some of the suppressors identified are required for epidermal wound closure, another Pvr-mediated wound response, some are required for hemocyte spreading *in vitro*, and some are required for both. One of the downstream factors, Mask, is also required for efficient wound-induced hemocyte spreading *in vivo*. Our data reveals that Pvr signaling is required for wound responses in hemocytes (cell spreading) and defines distinct downstream signaling factors that are required for either epidermal wound closure or hemocyte spreading.

## Introduction

*Drosophila* larvae have emerged as a useful system to study tissue repair responses (Tsai et al. 2018), including wound closure (WC) (Baek et al. 2010; Galko and Krasnow 2004; Kakanj et al. 2016), epidermal cell-cell fusion (Lee et al. 2017; Wang et al. 2015) and basement membrane dynamics (Ramos-Lewis et al. 2018). Following injury, larval barrier epithelial cells at the wound-edge locally detach from the apical cuticle and migrate into the wound gap. This process requires both JNK signaling (Galko and Krasnow 2004; Lee et al. 2019; Lesch et al. 2010) and Pvr signaling (Wu et al. 2009). The latter is required in some manner for epithelial extension into the wound site, though it has been difficult to identify downstream genes of this pathway given a lack of pathway reporters that function well *in vivo* during the larval stage.

*Drosophila* is also a good system for studying damage-induced inflammatory responses (Brock et al. 2008; Stramer and Dionne 2014). Hemocyte responses to wounding in *Drosophila* are remarkably stage-specific. The recruitment of hemocytes to wounds during the non-locomotory embryonic (Stramer et al. 2005) and pupal stages (Moreira et al. 2011) is primarily through directed cell migration of hemocytes. These migrations require hydrogen peroxide (Moreira et al. 2010) and likely other cues (Weavers et al. 2016). Larvae, which are a locomotory foraging stage that follows embryogenesis and precedes pupariation and, have a different mechanism of recruiting hemocytes to damaged tissue. In larvae, circulating hemocytes patrol the open body cavity and adhere to damaged tissue if they encounter it (Babcock et al. 2008). Once at the wound, attached hemocytes spread, change from an approximately spherical to a flattened fan-like morphology, and phagocytose cell debris. At the larval stage, even hemocytes close to the wound do not respond to it through directed migration (Babcock et al. 2008). Some hints about the molecules required for blood cell attachment have been gleaned from other insect species (Levin et al. 2005; Nardi et al. 2006) and from vertebrates (Eming et al. 2007). Likewise, some studies of *Drosophila* cell morphology have been performed in hemocyte-like cells *in vitro* (D’Ambrosio and Vale 2010; Kiger et al. 2003) and even in response to wounding *in vivo* (Kadandale et al. 2010). However, the molecules required for wound-induced spreading *in vivo* and their relationship to *in vitro* observations remain unclear.

Pvr is a *Drosophila* receptor tyrosine kinase (RTK) related to the vertebrate VEGF receptor (Cho et al. 2002; Heino et al. 2001). Pvr controls a variety of developmental signaling events including hemocyte differentiation (Mondal et al. 2014), migration (Cho et al. 2002; Wood et al. 2006), and survival (Bruckner et al. 2004; Munier et al. 2002; Zettervall et al. 2004). Pvr is also required for epithelial developmental migrations (Garlena et al. 2015; Harris et al. 2007; Ishimaru et al. 2004; McDonald et al. 2003) and for epidermal WC at the larval stage (Wu et al. 2009). Because Pvr is an RTK it presumably connects to a fairly canonical RTK signaling pathway downstream and some studies have identified downstream players in certain contexts (Fernández-Espartero et al. 2013; Jékely et al. 2005; McDonald et al. 2003). Notably, however, reliable reporters for monitoring pathway activity *in vivo* have been difficult to come by for this pathway. An alternative approach to finding pathway components, one with prior success for analyzing RTK pathways is genetic modifier screening (Smith et al. 2002; Sullivan and Rubin 2002). Here, we took advantage of the lethality of Pvr overexpression in the larval epidermis (Wu et al. 2009) to design a suppressor screen that could, in theory, identify downstream signaling components in this tissue. We then cross-checked the suppressors identified by the screen to see if they were required for larval epidermal WC or for hemocyte spreading at wound sites. This strategy revealed both shared and distinct downstream components for Pvr signaling in mediating epidermal WC and hemocyte spreading.

## Materials and Methods

### Genetics

*Drosophila* were reared on standard cornmeal medium under a 12 h light-dark cycle. All crosses were cultured at 25 °C unless indicated. *w^1118^* was used as a control strain. *Pvr^c02859^* is a hypomorphic allele (Cho et al. 2002; Wu et al. 2009). *Pvr^MI04181^* (Venken et al. 2011), referred to as *Pvr^null^*, contains a splice acceptor and a stop cassette in an early Pvr intron which leads to truncation. *Pvf1^EP1624^*, here referred to as *Pvf1^null^*, is a null allele (Cho et al. 2002; Wu et al. 2009). *Pvf2^c06947^*, here referred to as *Pvf2^hypo^*, is a hypomorphic allele (CHO *et al*. 2002). *Pvf3^M044168^*, here referred to as *Pvf3^null^*, contains a splice acceptor and a stop cassette in an early Pvf3 intron which leads to truncation (Venken et al. 2011).

The GAL4/UAS system was used to drive tissue-specific gene expression of transgenes under UAS control (Brand and Perrimon 1993). For larval hemocytes, *hmlΔ-Gal4* was used (Sinenko and Mathey-Prevot 2004); for the embryonic and larval epidermis, *e22c-Gal4* was used (Lawrence et al. 1995); for the larval epidermis, *A58-Gal4* was used (Galko and Krasnow 2004). To increase Pvr expression or activation in specific tissues, various Gal4 drivers were crossed to either *UAS-Pvr* or *UAS-λPvr* (Duchek et al. 2001). For the hemocyte spreading assay, we used *hmlΔ-Gal4*, *UAS-GFP or hmlΔ-Gal4* (Sinenko and Mathey-Prevot 2004), *UAS-lifeact-GFP* (Hatan et al. 2011). For visualizing WC, we used *e22c-Gal4*, *UAS-src-GFP*, *UAS-DsRed2-Nuc* or *A58-Gal4*, *UAS-src-GFP*, *UAS-DsRed2Nuc* (Lesch et al. 2010). *e22c-Gal4*, *UAS-src-GFP*, *UAS-DsRed2Nuc*; *tubP-gal80^ts^* was used where temporal control of the Gal4/UAS system was needed (McGuire et al. 2004).

*UAS-RNAi* lines employed were: From Vienna Drosophila Research Center (VDRC) (Dietzl et al. 2007): *KK108550* (*MKK3^RNAi#1^*), *GD7546* (*MKK3^RNAi#2^*), *KK100471* (*CG1227^RNAi^*), *GD14375* (*Pvr_RNAi#1_*). Note: lines are listed as-construct ID (GeneX^RNAi^). *UAS-RNAi* lines from the TRiP Bloomington collection (Ni et al. 2011) were: *JF01355* (*Luciferase^RNAi^*), *JF02478* (*Ras^RNAi#2^*), *HMS01294* (*Ras^RNAi#3^*), *HMS01979* (*Vav^RNAi^*), *HMS00173* (*Erk^RNAi^*), *HMS05002* (*MKK3^RNAi#3^*), *JF02770* (*PI3K92E^RNAi^*), *HMS00007* (*Akt^RNAi^*), *GL00156* (*Tor^RNAi#1^*), *HMS00904* (*Tor^RNAi#2^*), *JF02717* (*drk^RNAi^*), *HMS01045* (*mask^RNAi^*), *JF01792* (*Ck1α^RNAi#1^*), *GL0021* (*Ck1α^RNAi#2^*), *GL00250* (*GckIII^RNAi^*). *UAS-RNAi* lines from NIG-Fly (http://www.shigen.nig.ac.jp/fly/nigfly/index.jsp) were: *9375R-1* (*Ras85D^RNAi^*), *7717R-1* (*MEKK1^RNAi^*), *1587R-1* (*Crk^RNAi^*), *6313R-2* (*mask^RNAi#2^*), *8222R-3* (*Pvr^RNAi#2^*).

Other transgenic lines from Bloomington Stock Center: #9490, *w*\*; *TM6B*, *P*{*w*[+*mC*]=*tubP-GAL80*}*OV3*, *Tb^1^/TM^3^*, *Sb^1^* (Balancer Stock containing Gal80). #8529, *w*\*; *P*{*w*[+*mC*]=*UAS-lacZ.Exel*}*2* (used as UAS control). #64196, *w*; P{UAS-Ras85D.V12}2* (constitutively active form of Ras85D)(Lee et al. 1996). #19989, *P{y[+t7.7]=Mae-UAS.6.11}lic[GG01785]/FM7c* (overexpresses MKK3) (Beinert et al. 2004). #59005, *P{UAS-p38b.DN}1* (dominant negative form of *p38b*) (Adachi-Yamada et al. 1999). #5788, *P{UAS-Ras85D.K}5-1* (wild type Ras85D) (Karim and Rubin 1998). #4845, *P{UAS-Ras85D.N17}TL1* (dominate negative form of Ras85D) (Lee et al. 1996). #30139, *w[1118]; P{w[+mC]=Hml-GAL4.Delta}2 (hmlΔ-Gal4*). #30140, *w[1118]; P{w[+mC]=Hml-GAL4.Delta}2, P{w[+mC]=UAS-2xEGFP}AH2 (hmlΔ-Gal4, UAS-GFP*) (Sinenko and Mathey-Prevot 2004). #35544, *y[1] w[*]; P{y[+t*] w[+mC]=UAS-Lifeact-GFP}VIE-260B (UAS-lifeact-GFP*) (Hatan et al. 2011).

### Scanning Electron Microscopy (SEM)

Dissected larval epidermis were fixed in 3% glutaraldehyde/2% paraformaldehyde with 2.5% DMSO in 0.2 M sodium phosphate buffer for 15 min. Samples were then dehydrated in graded ethanol concentrations and hexamethyldisilazane. Next, processed samples were mounted on to double-stick carbon tabs (Ted Pella. Inc., Redding, CA), which have been previously mounted on to glass microscope slides. The samples were then coated under vacuum using a Balzer MED 010 evaporator (Technotrade International, Manchester, NH) with platinum alloy for a thickness of 25 nm, then immediately flash carbon coated under vacuum. The samples were transferred to a desiccator for examination at a later date. Samples were examined/imaged in a JSM-5910 scanning electron microscope (JEOL, USA, Inc., Peabody, MA) at an accelerating voltage of 5 kV. To quantify the SEM results, three to five (350X) images of each wound and three to twelve animals for each genotype were collected. These images were given to four or more persons to blindly score the hemocyte spreading phenotype. Percentages of hemocytes at the wound sites showing spreading morphology were binned into 0%, 25%, 50%, 75% and 100%. Scoring results of each image from different persons were averaged. Multiple images from the same animals were then averaged to obtain a “spreading index”.

### Pvf1 enrichment

The plasmid containing Pvf1d was transformed into BL21DE3 *E. coli* cells for overexpression. Cells were grown in Luria-Bertani broth at 37°C to an A600 density of 0.6 and Pvf1d (truncated version of Pvf1 containing only the VEGF-like domain) overexpression was achieved by induction with 1 mM Isopropyl ß-D-ThioGalactoside (IPTG) for 3 hours. Cell pellets were harvested and resuspended in lysis buffer containing 20 mM Tris pH8.0, 0.1 M NaCl, 5% glycerol, 1 mM Ethylene diamine tetra-acetic acid (EDTA), and 0.1 M Dithiothreitol (DTT). Cells were lysed using a French press and the inclusion bodies containing the overexpressed Pvf1d were collected by centrifugation at 15000 rpm for 30 minutes. Inclusion bodies were washed with the lysis buffer and stored in aliquots at −80°C. Approximately 1 gram of inclusion body was resuspended in lysis buffer containing 8 M Urea and dialysed overnight against the same buffer. Refolding of Pvf1d was achieved by overnight dialysis against buffer containing 50 mM N-cyclohexyl-3-aminopropanesulfonic acid (CAPS) pH 10.5, 50 mM NaCl, 5% glycerol, and 5 mM cysteine. Prior to dialysis, protein concentration was adjusted to 0.2 mg/ml and the dialysis step was repeated two more times. Subsequently, protein was cleared of precipitates by centrifugation and purified into a storage buffer containing 20 mM CAPS pH 10.5, 50 mM NaCl and 2.5% glycerol by size exclusion chromatography.

### *In vitro* hemocyte spreading assay

Hemocytes were isolated from wandering third instar larvae (genotype: *w*;*hmlΔ-Gal4*,*UAS-GFP +/-UAS-RNAi transgene*) using a protocol modified from (Kadandale et al. 2010).

Approximately 150 mg of larvae (~ 100) were collected into a cell strainer (70 μm pore size) and washed once in Phosphate Buffered Saline (PBS). The rinsed larvae were crushed within the cell strainer in a 35 mm sterile cell culture dish with the cap-end of an eppendorf tube. The crushate containing hemocytes was filtered into the 35 mm dish by washing the crushed larvae twice with 500 μl of PBS. The hemocyte-containing filtrate was collected into a 1.5 ml eppendorf tube and was centrifuged for 1 min at 1000 rpm to remove particulates. The supernatant was re-centrifuged at 2000 rpm for 2 min to collect hemocytes. The hemocyte-containing pellet was resuspended in 500 μl of room temperature Schneider’s *Drosophila* culture medium (GIBCO, Invitrogen). ~1 × 10^5^ cells suspended in the culture media described above were plated onto coverslips (Corning) that were placed in a sterile 24 well culture well (Corning). After plating, 1 μl of 44 ng/μl recombinant Pvf1 protein was added to the culture and the cells were treated for 1 hr at 25 °C. 1 μl of 1X PBS was added to control cells instead of Pvf1 and were cultured for 1 hr at 25 °C. Phalloidin staining: After 1 hr of Pvf1 or control treatment, the cells were washed once with PBS and fixed for ten min with 4% paraformaldehyde before washing three times with PBS. The cells were permeabilized with 0.1% Triton X-100 (TX-100) in PBS (washing buffer) for 10 min and then incubated in blocking buffer (3% BSA, 0.1% TX-100 prepared in 1X PBS) for 30 min at room temperature. The cells were stained overnight at 4 °C with a 1:50 dilution of phalloidin-Alexa 546 (Invitrogen) made in blocking buffer followed by three washes each of 5 min. After washing, the coverslips containing phalloidin-stained cells were lifted off the well and mounted on to a glass slide (Fisher Scientific) using a drop (around 3 μl) of mounting media (Vectashield, Vector Laboratories). The coverslips were sealed to the glass slide with clear nail polish and stored at 4 °C until imaged. Anti-Phospho-Pvr antibody staining: Phospho-Pvr (pPvr) antibody (monoclonal antibody) that detects the phosphorylation of Pvr at Tyr 1426 (Janssens et al. 2010) was a generous gift from Dr. P. Rørth (Institute of Molecular and Cell Biology, Proteos, Singapore). Hemocytes were isolated and processed as mentioned above until the completion of blocking. Staining was performed with a 1:5 dilution of anti-pPvr (diluted in blocking buffer) at 4 °C overnight. The secondary antibody was Goat anti-mouse DyLight 649 (Jackson ImmunoResearch Laboratories) which was bound for 1 hr at room temperature before washing and mounting onto glass slides as described above.

### Hemocyte spreading screen

A more streamlined version of the above spreading assay was developed for the purposes of screening. In this protocol, select *UAS-RNAi* lines were crossed with the screening stock (*hmlΔ-Gal4*, *UAS-lifeact-GFP*, *UAS-λPvr*) at 25°C. For each cross, ~10 mid-3^rd^ instar larvae (5 days after egg lay) carrying the Gal4 driver, *UAS-lifeact-GFP*, *UAS-λPvr* (*UAS-Pvr^CA^*) and the candidate *UAS-RNAi* transgene were selected and placed in a glass dissection well containing PBS. Larvae were washed with 70% ethanol and PBS and then briefly kept in 300 μl of PBS. Hemocytes were released from the larvae by nicking their posterior ends with dissection scissors (Fine Science Tools, #15000-02). Collected hemocytes were transferred to an ice-cold low-retention tube (Fisher, #02-681-331). Collected hemocytes were seeded into an 8-well chamber slide (Millipore, PEZGS0816) and allowed to spread for one hour at room temperature. After spreading, samples were fixed with 3.7% formaldehyde for five minutes, washed with PBS, and mounted in Vectashield before imaging with Olympus FV1000 Confocal microscope with Fluoview software and 60x oil lens. ImageJ was used to manually measure the longest axis of individual hemocytes. Overlapping hemocytes were excluded from measurement to avoid potential interference between cells. To measure the hemocyte size before spreading, hemocytes were fixed (as above) right after isolation and washed with PBS before resuspending in Vectashield and mounting onto slides for imaging.

### Lethality suppressor screen

Candidate *UAS-RNAi* lines were crossed with the screening stock (*UAS-Pvr*; *A58-Gal4/TM6B*, *tubP-Gal80*) at 22.5°C, at which the best signal to noise ratio of the screen was observed. Flies were transferred onto fresh vials every two days. *UAS-Luciferase^RNAi^* and *UAS-Pvr^RNAi#2^* were used as negative and positive controls, respectively. Larvae, pupae, and/or adults emerging from the different crosses were observed six to nine days after egg laying. The *UAS-Luciferase^RNAi^* control group does not survive to the prepupal and pupal stages, whereas the *UAS-Pvr^RNAi#2^* group survives until adult stage. Candidate genes were scored as putative suppressors when their corresponding *UAS-RNAi* transgenes delayed the lethal stage to prepupae or pupae. Median suppression was defined by the observation of three to five pupae/prepupae in a single vial (annotated in Figure 3C with “+”). Strong suppression was defined by the observation of six or more pupae/prepupae in a single vial (annotated in Figure 3C with “++”). No suppression “–” or variable suppression across multiple trials “+/-” were annotated in Figure 3C.

**Figure 1.**
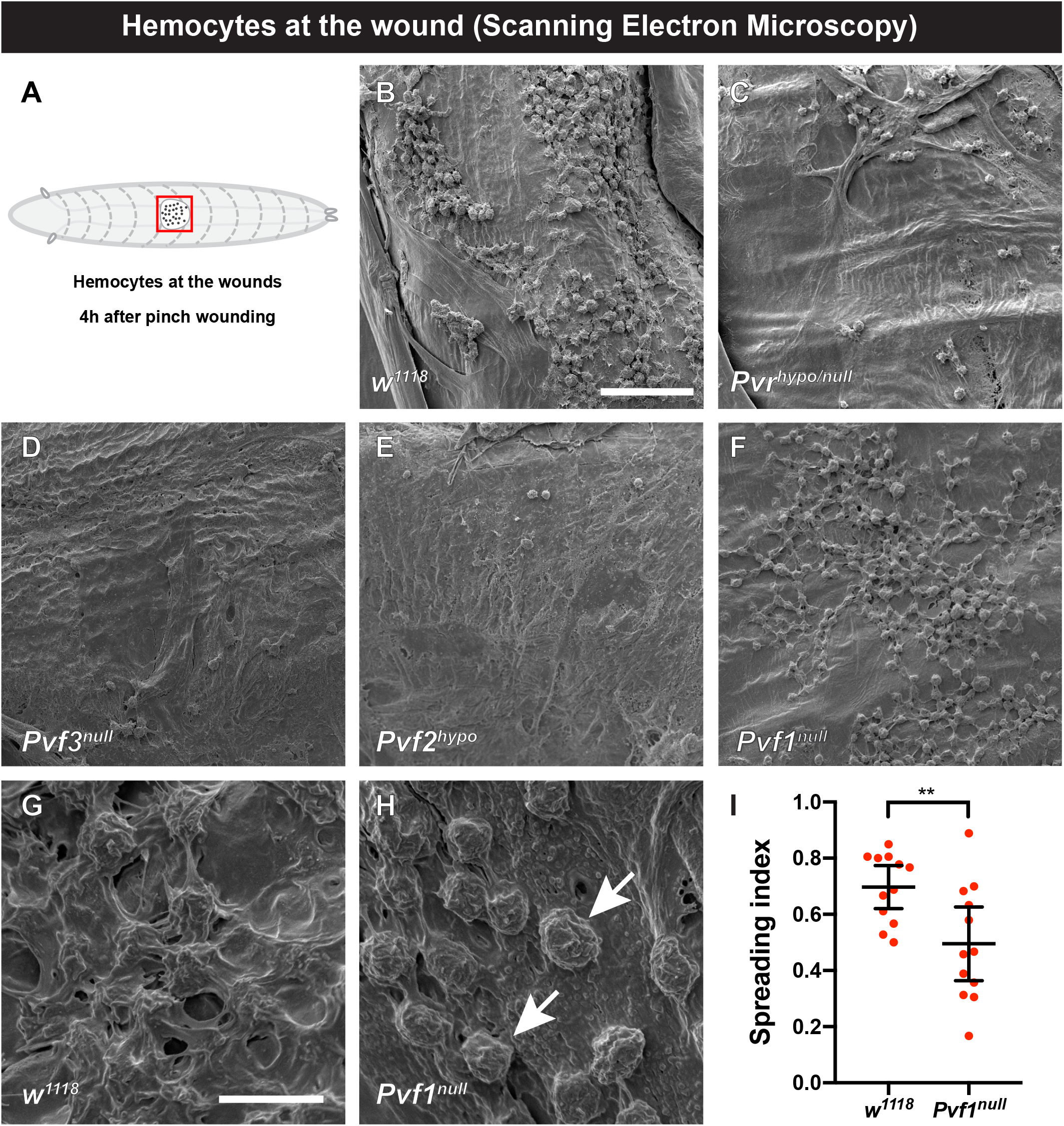
Pvr and Pvf1 are required for hemoctye spreading at larval wound sites. (A) Cartoon of third instar *Drosophila* larva (anterior to left, posterior to right) red square highlighting the region of interest (clear oval, the wound, and black dots, hemocytes) for scanning electron microscopy (SEM) analysis of pinch wounds. (B-H) Scanning electron micrographs of wounded and dissected third instar larvae of the indicated genotypes to visualize wound-adherent blood cells. (B) *w^1118^* control (C) *Pv^nullo/hypol^* (D) *Pvf3^null^* (E) *Pvf2^hypo^* (F) *Pvf1^null^* Scale bar in (B) = 50 μm and applies to (B-F) (G) Close-up of spread hemocytes, *w^1118^*. (H) Close-up of unspread hemocytes indicated by arrows, *Pvf1^null^*. Scale bar in (G) = 10 μm and applies to (G-H). (I) Quantitation of blood cell spreading in control larvae versus *Pvf1^null^* mutant larvae. n = 12. Data are mean with 95% CI. \*\**P*<0.01 (unpaired two-tailed *t*-test).

**Figure 2.**
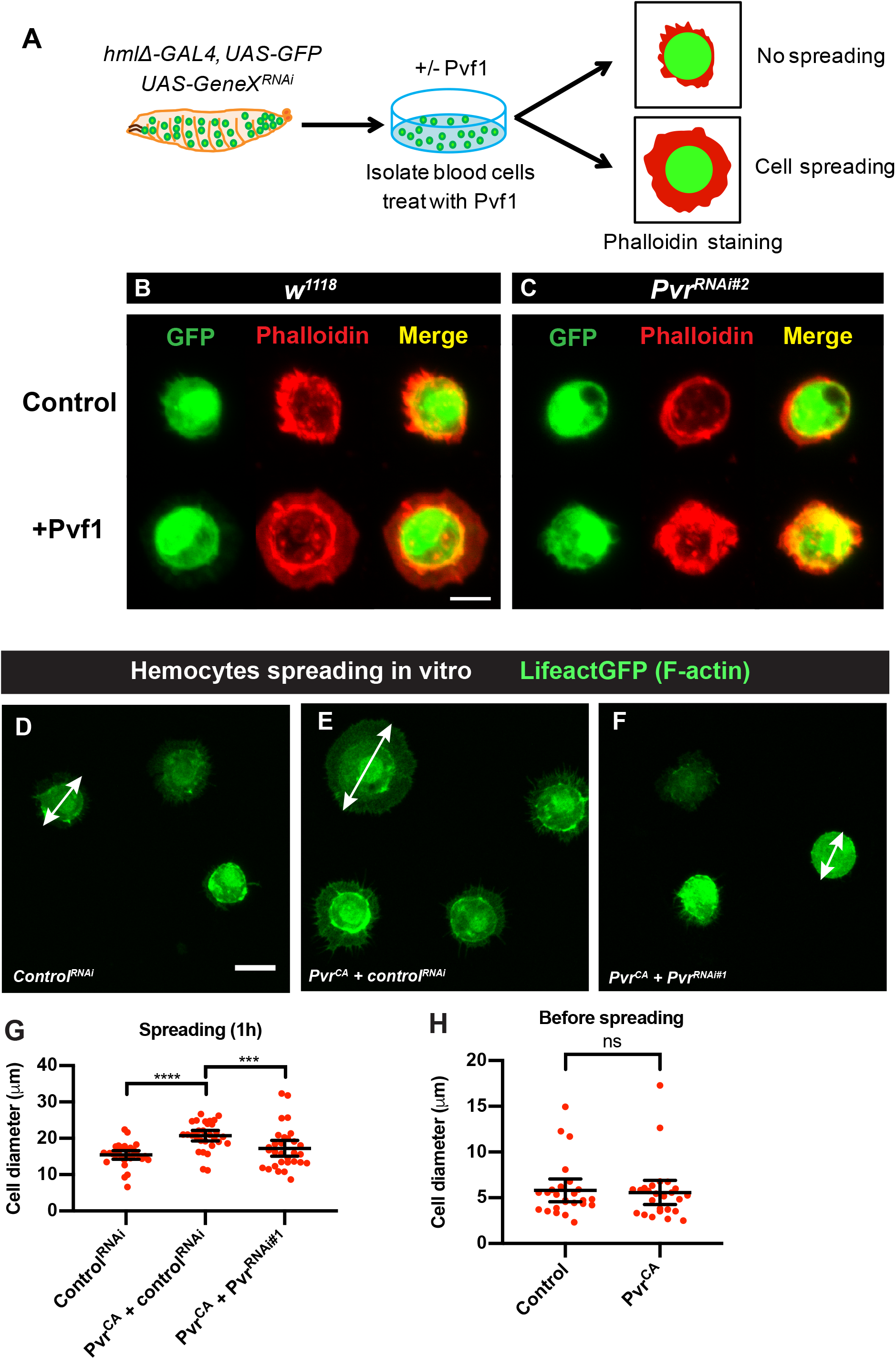
Testing the role of Pvr/Pvf1 in hemocyte spreading with *in vitro* assays. (A) Schematic of hemocyte spreading assay for treatment with Pvf1. (B) Untreated control hemocytes (*w^1118^*; *hemolectinΔ-Gal4*, *UAS-GFP*) and treated (+ enriched Pvf1 protein) hemocytes are shown. Blood cells were harvested from larvae, plated *in vitro*, fixed, and visualized with the GFP lineage label (green, left column), phalloidin to label filamentous actin (red, middle column), or both (merge, right column) in the absence (top row) or presence (bottom row) of enriched Pvf1 protein. Scale bar in (B) = 10 μm and applies to (B-C). (C) Same experiment as in (B) but now the hemocytes are also expressing a *UAS-Pvr^RNAi^* transgene (bottom row) or not (top row) to test whether the spreading response observed upon addition of Pvf1 protein depends on functional Pvr expression. (D-F) Morphology of plated hemocytes (*w^1118^*; *hemolectinΔGal4*, *UAS-LifeActGFP*, green) with the indicated transgenes. Scale bar in (D) = 10 μm and applies (D-F). Double-headed Arrow in (D-F) are examples of cell longest diameters. (D) *UAS-Control^RNAi^*. (E) *UAS-Pvr^CA^ + UAS-Control^RNAi^*. (F) *UAS-Pvr^CA^* + *UAS-Pvr^RNAi#1^*. (G-H) Quantitation of hemocyte cell diameters (μm) of the indicated genotypes after 1 hour of plating (G) or before plating (H) to test whether expression of *UAS-Pvr^CA^* affects hemocyte size in any way. (G,H) Each dot represents the diameter of a single cell. Error bars: mean with 95% CI. (G) n = 30, (Kruskal-Wallis multiple comparisons test). (H) n = 25; ns, not significant (Kolmogorov-Smirnov test).

**Figure 3.**
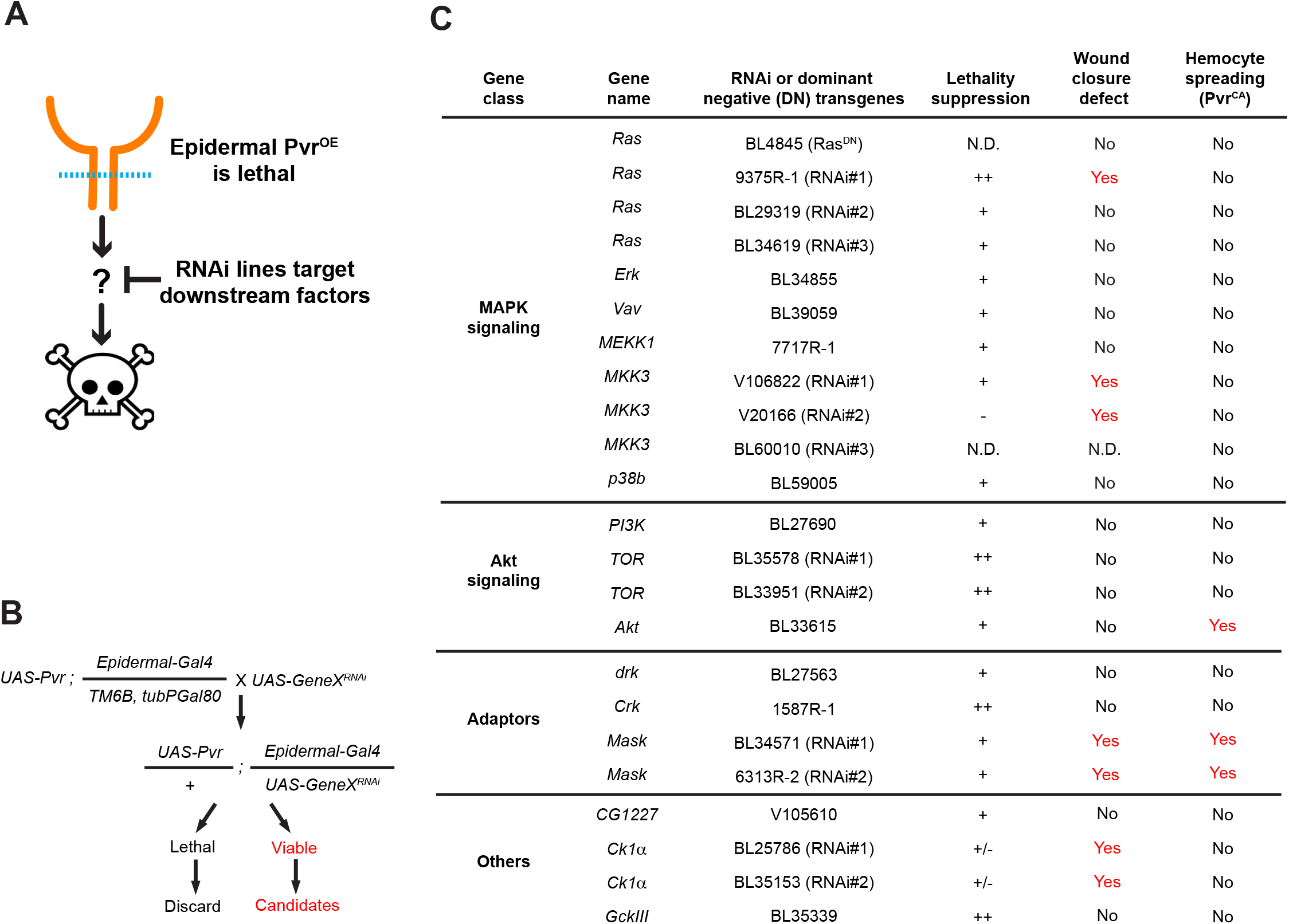
Targeted genetic screen for suppressors of Pvr-induced lethality. (A) Conceptual schematic of genetic screen. Pvr overexpression in the larval epidermis is lethal. We screened for RNAi lines (targeting possible/probable downstream components of RTK signaling) that, when co-expressed with Pvr, could suppress this lethality (B) Genetic scheme of the screen, illustrating the genotypes, crosses, and scoring involved. (C) Lethality suppressors from the screen, organized by gene class. Also shown are whether the suppressors affected epidermal wound closure (WC) at the larval stage and/or hemocyte spreading in the *in vitro* assay (see Fig. 2). For the strength of lethality suppressions: ++, strong suppression. +, median suppression. -, no suppression. +/-, variable suppression effects. N.D., not determined.

### Larval wound closure assay

Pinch wounding of the larvae was carried out according to our detailed protocol (Burra et al. 2013). In cases where early expression of a UAS transgene was lethal (*UAS-Akt^RNAi^*), larvae bearing *tub-gal80^ts^*, the Gal4 driver and toxic UAS transgene were raised for six days at 18 °C to begin development, shifted to 32 °C for two days to reach mid-third-instar, and then allowed to recover at 25 °C following pinch wounding. Pinch wounds were scored as “open” if the initial wound gap remained after 24 hours, and as “closed” if a continuous epidermal sheet was observed at the wound site. To calculate the percentage of larvae with open wounds, three sets of N ≥ 8 per genotype were pinched and scored for open wounds under a fluorescent stereo microscope (Leica MZ16FA with Planapo 1.6x objective and appropriate filters). To further examine wound morphology, the third instar larval epidermis was dissected and processed as detailed previously (Burra et al. 2013). To highlight epidermal morphology, a mouse monoclonal antibody against Fasciclin III was used (1:50; Developmental Studies Hybridoma Bank). An Olympus FV1000 Confocal microscope, Olympus 20x oil lens and Fluoview software were used to obtain images of the dissected epidermal whole mounts.

### Statistical analyses

For statistical analysis of the WC phenotype between genotypes, one-way ANOVA (Dunn’s multiple comparisons) were used to test the significance of experiments.

For statistical analysis of hemocyte spreading, if the data of all the genotypes passed D’Agostino and Pearson omnibus normality test, unpaired two-tailed *t*-test (two groups) or one-way ANOVA (more than two group, Dunn’s multiple comparisons) were used to test the significance of experiments. When data from one or more genotypes did not pass D’Agostino and Pearson ominbus normality test, Kolmogorov-Smirnov test (two groups) or Kruskal-Wallis test (more than two groups, Dunn’s multiple comparisons) were used to test the significance of experiments. For all quantitations: ns, not significant; \**P*<0.05; \*\**P*<0.01; \*\*\**P*<0.001; \*\*\*\**P*<0.0001.

#### Data availability

Strains and plasmids are available upon request. A supplemental material file in the online of this article contains Figure S1 and Table S1 (genotypes used in each figure). Supplemental material available at FigShare.

## Results

### Pvr and Pvf1 are required for hemocyte spreading at wound sites

In *Drosophila* larvae, circulating hemocytes adhere to wound sites if they encounter the wound surface by chance (Babcock et al. 2008). Once there, they assume a spread morphology and phagocytose wound-associated debris (Babcock et al. 2008). We sought to identify factors that might be responsible for hemoctye spreading *in vivo*. We began our search with transmembrane proteins known to be expressed on hemocytes and known to affect hemocyte biology. Pvr (PDGF/VEGF-related receptor) fits these criteria (Bruckner et al. 2004; Cho et al. 2002; Heino et al. 2001). To observe hemocytes at wound sites, we pinch-wounded (Burra et al. 2013) third instar *Drosophila* larvae and used scanning electron microscopy (SEM) to examine the morphology of wound-adherent hemocytes (see schematic in Fig. 1A). In control larvae (Fig. 1B-see also Table S1 for list of genotypes relevant to each figure panel) large numbers of hemocytes bound to the wound and assumed a spread morphology. In *Pvr^null/hypo^* (see materials and methods and Table S1 for allele designations) there were much fewer hemocytes at the wound site (Fig. 1C). This is to be expected, as Pvr is required for hemocyte survival in embryos (Bruckner et al. 2004). Further, Pvr activation (Zettervall et al. 2004) or Pvf2 overexpression (Munier et al. 2002) can drive hemocyte proliferation at the larval stage. We also observed greatly reduced hemocyte numbers in *Pvf2^hypo^* and in *Pvf3^null^* mutants at the wound sites (Fig. 1D-E), suggesting that these ligands may also be required for hemocyte survival. The third VEGF-like ligand, Pvf1, showed a different phenotype at wound sites (Fig. 1F) compared to *Pvf2* and *Pvf3* mutants. While hemocytes were present in substantial numbers at wound sites within *Pvf1^null^* larvae, closer examination revealed that they possessed a morphology distinct from controls. Higher magnification views of control larvae (Fig. 1G) show that spread hemocytes formed a dense and interlinked network of cell processes over the wound site. In *Pvf1^null^* mutants the hemocytes adhered, but had a distinctly rounded morphology, with few broad and flattened membrane sheets, even when in close proximity to each other (Fig. 1H). Quantitation of the spreading index (see materials and methods) between these two genotypes revealed a significant difference in visible morphology (Fig. 1I). In sum, Pvr and two of its ligands, Pvf2 and Pvf3, are required for normal numbers of wound-adherent hemocytes, while Pvf1 is required for these cells to assume a spread morphology at the wound site.

### *In vitro* assays for hemocyte spreading-a flattened lamellar morphology induced by Pvf1 application or Pvr activation

*In vivo* loss of function analysis suggested that Pvf1, possibly through the Pvr receptor, is required for hemocyte spreading. We tested this in another way, by modifying an *in vitro* assay for hemocyte spreading (Fig. 2A) (D’Ambrosio and Vale 2010; Kiger et al. 2003). Lineage-labeled plasmatocytes (*hemolectinΔ-Gal4*, *UAS-GFP*) were collected from third instar larvae, plated, and exposed to enriched (Fig. S1A) Pvf1 VEGF-like domain (see methods). This enriched protein was active, as assessed by its ability to cause Pvr phosphorylation in isolated hemocytes (Fig. S1B-B’). The phosphorylation signal was specific, as it depended upon Pvr expression in the isolated hemocytes (Fig. S1C-C’).

Control hemocytes plated *in vitro* assumed a rounded morphology, as assessed by the cytoplasmic GFP label (Fig. 2B). When stained with phalloidin, which labels filamentous actin, these cells exhibited a peripheral ring of dense actin filaments (Fig. 2B, top row). Exposure to enriched and active Pvf1 VEGF-like domain during the period of plating altered the morphology of these cells-they now exhibited a large lamellipodial-like fan extending outwards from the peripheral actin ring (Fig. 2B, bottom row). To determine whether this *in vitro* Pvf1-dependent spreading requires the Pvr receptor, we isolated hemocytes co-expressing a *UAS-Pvr^RNAi^* transgene whose efficacy has been verified in other assays (Lopez-Bellido et al. 2019; Wu et al. 2009). In the absence of exogenous Pvf1 protein, hemocytes expressing *UAS-Pvr^RNAi#2^* had a morphology and actin distribution similar to controls (Fig. 2C, top row). These same cells, when plated in the presence of Pvf1 protein, exhibited an apparent increase in cellular actin staining but did not spread outwards to form a lamellipodial fan (Fig. 2C, bottom row).

Finally, we determined whether hyperactivation of Pvr *in vivo* (through expression of the constitutively active *UAS-Pvr^CA^* transgene (Duchek et al. 2001) could directly lead to spreading of hemocytes. Hemocytes expressing a *UAS-LifeactGFP* transgene (to label filamentous actin) and a *UAS-control^RNAi^* transgene (Fig. 2D) possessed a simple rounded morphology *in vitro*. By contrast, hemocytes co-expressing *UAS-Pvr^CA^* and *UAS-Luciferase^RNAi^* transgene (to equalize the number of UAS transgenes in the experimental setup) exhibited prominent lamellipodial fans (Fig. 2E) similar to those observed upon co-culture with the Pvf1 VEGF-like domain (Fig. 2B, bottom row). The spreading phenotype of different genotypes was measured based on the average of individual cell diameters measured at the longest axis for each cell. Cell diameters of hemocytes expressing *UAS-Pvr^CA^* and *UAS-control^RNAi^* were significantly larger than control (Fig. 2G). The presence of *UAS-Pvr^CA^–induced* lamellipodial fans was dependent upon Pvr, as co-expression of *UAS-Pvr^CA^* and *UAS-Pvr^RNAi#1^* led to hemocytes with a simple rounded morphology (Fig. 2F,G). The fan-like morphology of hemocytes expressing activated Pvr was not simply due to an increase in the original size of the hemocytes. When we measured cell size before plating (Fig. 2H), there was no difference in the cell diameter of hemocytes expressing *UAS-Pvr^CA^* versus controls. By contrast, *UAS-Pvr^CA^* –expressing hemocytes were of significantly greater diameter one hour after plating, an effect that was dependent upon expression of Pvr (Fig. 2G). Together, these data demonstrate that Pvf1 causes hemocyte spreading via Pvr activation.

### A suppressor screen for genes that act downstream of Pvr signaling

Pvr signaling has a unique place in *Drosophila* tissue damage responses in that it is required in multiple tissues for diverse cellular responses. In the larval epithelium, Pvr is required for wound closure (WC) (Wu et al. 2009), in nociceptive sensory neurons for the perception of noxious mechanical stimuli (Lopez-Bellido et al. 2019), and in hemocytes for spreading at wound sites (Fig. 1). A challenge in studying this pathway has been that there are no broadly useful reporters of downstream pathway activity. The anti-phospho-Pvr antibody used in Fig. S1 is only useful on isolated cells and not for wholemount tissue stains (data not shown). Given these challenges, we designed a genetic screen to efficiently identify genes that act downstream of Pvr activation. To do this, we took advantage of the fact that overexpression of Pvr in the *Drosophila* larval epidermis is lethal (Wu et al. 2009). The screen itself is a lethality suppressor screen (see conceptual schematic in Fig. 3A). We reasoned that co-expression of *UAS-RNAi* transgenes targeting potential downstream genes would suppress the lethality induced by overexpression of Pvr. The screening stock(s) and crossing scheme for the screen is depicted in Fig. 3B and hinges on the use of the Gal80 system (Vef et al. 2006) to suppress expression of *UAS-Pvr* and keep the screening stock alive. The candidate set of *UAS-RNAi* lines included known kinases and adaptors that act downstream of RTKs as well as a broader set of such genes. The first phenotype screened was the presence of pupae in the vials co-expressing *UAS-Pvr* and the *UAS-GeneX^RNAi^* transgenes. In total, about 600 genes were screened and 15 lethality suppressors were obtained (Fig. 3C). Many of the basic components of mitogen-activated protein kinase (MAPK) and Akt signaling, as well as a subset of common RTK adaptors and other kinases scored positive as suppressors. Ultimately, all of the lethality suppressors (and further RNAi or dominant-negative transgenes targeting them) were also screened for phenotypes in larval WC and *in vitro* hemocyte spreading (Fig. 3C, right columns) and a subset of these phenotypes are shown in the ensuing figures below.

### New wound closure genes-Ras, MKK3, and Mask

In the ideal case, Pvr signaling architecture would be similar between Pvr-induced lethality and WC and most or all of the lethality suppressors would then score positive as genes required for larval WC. This was not in fact observed (see discussion section below for possible explanations). Only a specific subset of the lethality suppressors were also identified as WC genes. When *UAS-Luciferase^RNAi^* transgenes (negative control) are expressed in the larval epidermis, pinch wounds close (Fig. 4A). By contrast, when *UAS-Pvr^RNAi^* transgenes are expressed (Fig. 4B, positive control) pinch wounds remain open at 24 hr post-wounding. The open-wound phenotypes observed upon expression of *UAS-RNAi* transgenes targeting Ras, a small GTPase (Fig. 4C), Mask, an adaptor protein (Fig. 4D), and MKK3, a MAP kinase kinase (Fig. 4E) are shown in Fig. 4, as is quantitation of the prevalence of these phenotypes (Fig. 4F). The lethality suppressor screen, while not perfect, was nonetheless quite fruitful at expanding our collection of known WC genes beyond the JNK and actin pathways (Brock et al. 2012; Lesch et al. 2010). Other genes that scored positive in this screen (CK1α) were also found in an analysis of adherens junctions at larval wound sites (Tsai and Galko 2019).

**Figure 4.**
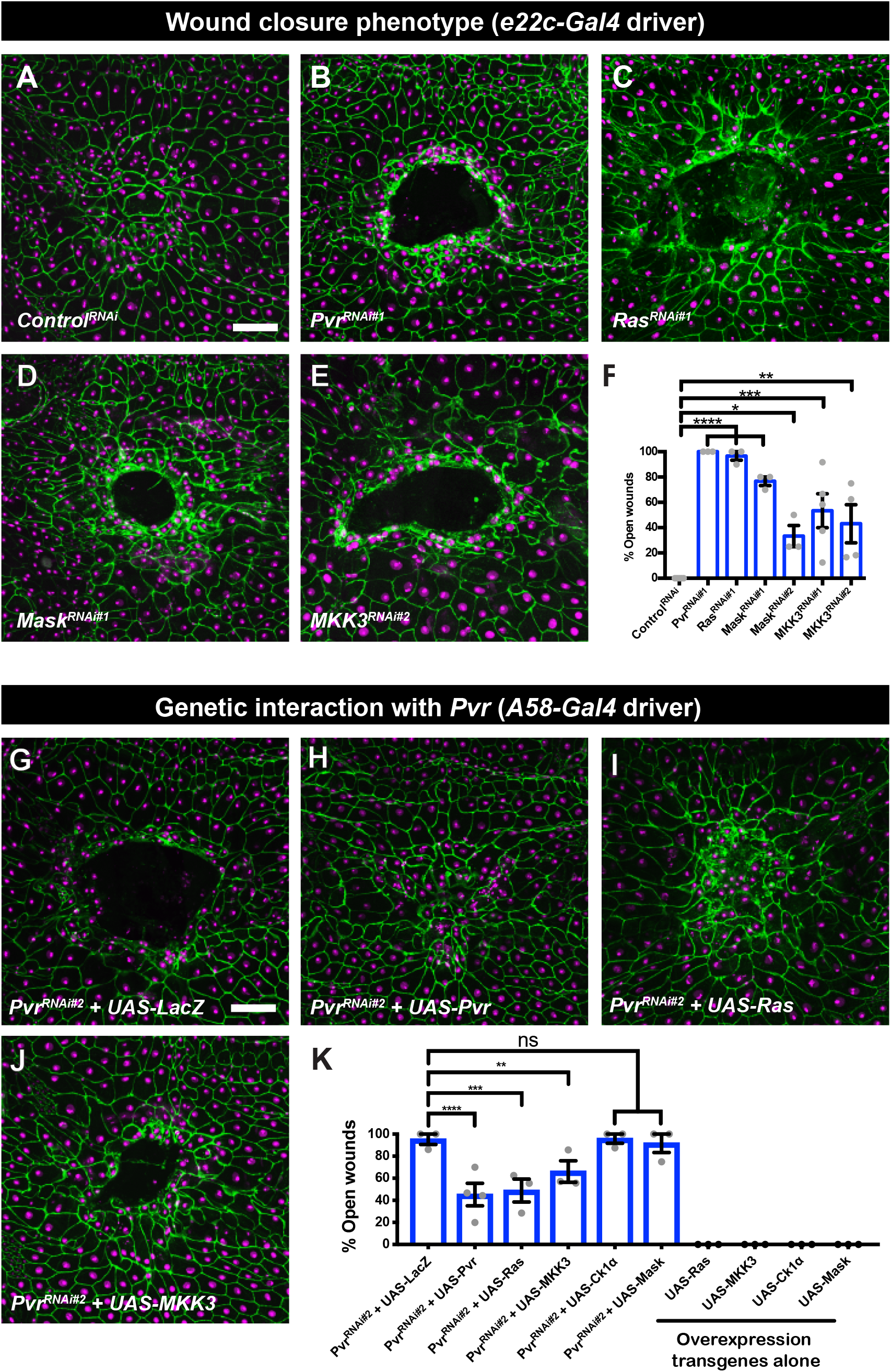
Epidermal wound closure phenotypes of select suppressors of Pvr-induced lethality and genetic interactions with Pvr. (A-E) Dissected epidermal whole mounts of wounded third instar larval epidermis, expressing *UAS-dsRed2Nuc* (Nuclei, magenta) *and UAS-src-GFP* (GFP, not shown) and expressing the indicated transgenes via *e22c-Gal4* driver, immunostained with anti-Fasciclin III (green). Open wounds appear as dark holes in the center. (A) *UAS-Control^RNAi^* (B) *UAS-Pvr^RNAi#1^* (C) *UAS-Pas^RNAi#1^* (D) *UAS-Mask^RNAi#1^* (E) *UAS-MKK3^RNAi#2^*. Scale bar, 50 μm in (A) is for (A-E). (F) Quantitation of larval WC phenotypes (% Open wounds) versus genotype. Each dot represents one set of n ≥ 8. Total three or more sets for each genotype. Error bar, mean ± S.E.M. One-way ANOVA with Dunn’s multiple comparisons. \*\*\*\**P*<0.0001, \*\*\**P*<0.001, \*\**P*<0.01, \**P*<0.05. (G-J) Epistasis. Ability of overexpression of select WC genes to rescue the WC phenotype of *UAS-Pvr^RNAi#2^*. Genotype of all panels: *w^1118^*; *UAS-Pvr^RNAi#2^*/+; *A58-Gal4*, *UAS-dsRed2Nuc*, *UAS-src-GFP* (not shown) plus the indicated overexpression transgene. (G) *UAS-Pvr^RNAi#2^* + *UAS-lacZ*. (H) *UAS-Pvr^RNAi#2^* + *UAS-Pvr*. (I) *UAS-Pvr^RNAi#2^* + *UAS-Ras*. (J) *UAS-Pvr^RNAi#2^* + *UAS-MKK3*. Scale bar, 50 μm in (G) is for (G-J). (K) Quantitation of epistasis experiments- % Open wounds versus the indicated genotypes. Each dot represents one set of n ≥ 8. Total three or more sets for each genotype. Error bar, mean ± S.E.M. One-way ANOVA multiple comparisons. \*\*\*\**P*<0.0001, \*\*\**P*<0.001, \*\**P*<0.01, ns, not significant.

Which, if any, of the identified WC genes act downstream of Pvr in the context of larval WC? We designed an experimental strategy (co-expression of *UAS-Pvr^RNA^* and a *UAS-cDNA* transgene for candidate genes) that would test this possibility. Certainly, suppression of the full WC defect caused by *UAS-Pvr^RNAi^* is a high bar, and might only be expected to be observed for those genes at or close to the top of the signaling pathway. Co-expression of an irrelevant gene (*UAS-LacZ*, negative control) was not capable of suppressing the open-wound phenotype observed upon expression of *UAS-Pvr^RNAi^* (Fig. 4G) indicating that titrating the Gal4/UAS system with a additional UAS sequences, by itself, was insufficient to suppress the WC phenotype. By contrast, co-expression of *UAS-Pvr* (positive control) suppressed the open wound phenotype of *UAS-Pvr^RNAi^* (Fig. 4H) about half of the time (Fig. 4K). Ras suppressed at a similar level (Fig. 4I, K) while MKK3 (Fig. 4J, K) was slightly weaker. Ck1α and Mask could not suppress (Fig. 4K). Of note, none of the *UAS-cDNA* overexpression transgenes caused an open wound phenotype on their own (Fig. 4K, right side). In sum, we have identified a number of new larval WC genes, some of which, by genetic epistasis, can be placed downstream of Pvr in this particular process.

### Mask and Akt act downstream of Pvr to mediate hemocyte spreading *in vitro*

We devised a parallel strategy to determine which of the Pvr lethality suppressors act downstream of Pvr in hemocyte spreading. This analysis was somewhat simpler, as we could ask whether each lethality suppressor could also suppress the hemocyte spreading induced by hemocyte expression of *UAS-Pvr^CA^* (see Fig. 5A schematic and Fig. 5B control). Co-expression of *UAS-RNAi* transgenes targeting either Akt (Fig. 5C,E) or Mask (Fig. 5D,F) resulted in a decrease of the expanded hemocyte cell diameter typically seen upon expression of *UAS-Pvr^CA^*. By contrast, *UAS-RNAi* and/or *UAS-DN* transgenes targeting MKK3 (Fig. 5G), Ck1α (Fig. 5H), or Ras (Fig. 5I) did not block *Pvr^CA^*-induced hemocyte spreading. Importantly, expressions of either *UAS-Akt^RNAi^* or *UAS-Mask^RNAi^* did not affect basal hemocyte spreading after one hour as measured by cell diameters (Fig. 5J). While some of the genes analyzed (in particular Ras, Ck1a, and MKK3) caused a general/baseline decrease in basal hemocyte spreading (Fig. 5J), Pvr-induced spreading was compared to the relevant baseline for each gene (Fig. 5E-I). These results demonstrate that some Pvr downstream factors (Mask) are shared between larval epidermal WC and hemocyte spreading while others, (Akt, Ck1a, MKK3, Ras) are specific for a particular cellular response.

**Figure 5.**
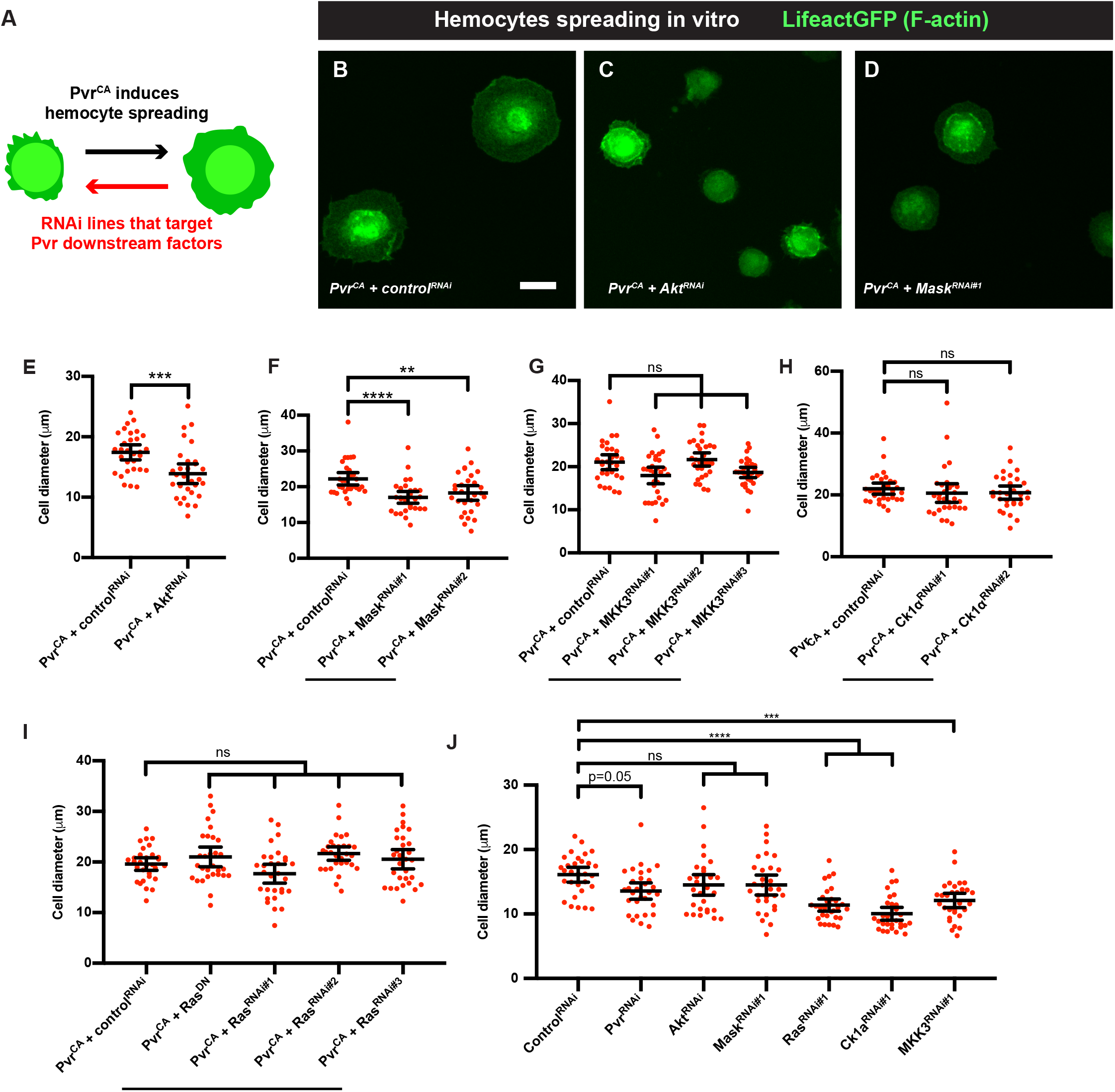
Hemocyte spreading phenotypes of select suppressors of Pvr-induced lethality. (A) Conceptual schematic of hemocyte spreading assay. Overexpression of a constitutive active form of Pvr (Pvr^CA^) in the larval hemocytes promote cell spreading. We screened for lethality suppressor RNAi lines that, when co-expressed with Pvr^CA^, could suppress the spreading phenotype. (B-D) Hemocyte morphology of hemocytes harvested from larvae of the genotype (*UAS-Pvr^CA^*, *hml Δ-Gal4*, *UAS-LifeactGFP*) plus the indicated transgenes, plated one hour *in vitro*, and visualized by the actin/lineage label (green). (B) *UAS-Pvr^CA^ + control^RNAi^*. (C) *UAS-Pvr^CA^ + UAS-Akt^RNAi^*. (D) *UAS-Pvr^CA^* + *UAS-Mask^RNAi#1^*. (E-J) Quantitation of hemocytes diameter (spreading) versus the indicated genotypes targeting particular genes. Each dot represents a single cell. n = 30. Error bars: mean with 95% CI. ****<0.0001. ***<P0.001. **P<0.01. ns, not significant. (E) Akt. Unpaired *t*-test. (F) Mask. Kruskal-Wallis multiple comparisons test. (G) MKK3. Kruskal-Wallis multiple comparisons test (H) Ck1α. Kruskal-Wallis multiple comparisons test (I) Ras. One-way ANOVA multiple comparison. (J) Expressions of *UAS-Akt_RNAi_* and *UAS-Mask^RNAi^* in hemocytes did not affect basal spreading, while *UAS-Ras^RNAi^*, *UAS-Ck1α^RNAi^* and *MKK3^RNAi^* reduced basal spreading. Kruskal-Wallis multiple comparisons test.

### Mask is also required for hemocyte spreading at wound sites *in vivo*

We next analyzed, using the SEM assay introduced in Figure 1, whether genes that have phenotypes in the *in vitro* hemocyte spreading assay (Mask and Akt) also affected wound-induced hemocyte spreading *in vivo*. As observed previously control hemocytes typically form a dense lawn on the wound surface (Fig. 6A) and, when analyzed at higher magnification, exhibit fan-like lamellipodial extensions either towards each other or towards the cuticle surface (Fig. 6D). In larvae expressing *UAS-Mask^RNAi^* in hemocytes, the wound-adherent cells appeared less dense (Fig. 6B) and possessed a wrinkled but rounded morphology that did not include lamellipodia extending either towards each other or the cuticular surface (Fig. 6E). In larvae expressing *UAS-Akt^RNAi^* in hemocytes (Fig. 6C) there appeared to be a survival defect similar to that observed for the Pvr, Pvf2 and Pvf3 ligands (Fig. 1C-E), as very few hemocytes were observed at the wound site. Quantitation of the spreading index in control versus *UAS-Mask^RNAi^*-expressing hemocytes (Fig. 6F) revealed a significant defect in spreading, indicating that for this gene, the *in vitro* spreading defect was an accurate predictor of a requirement for spreading *in vivo*.

**Figure 6.**
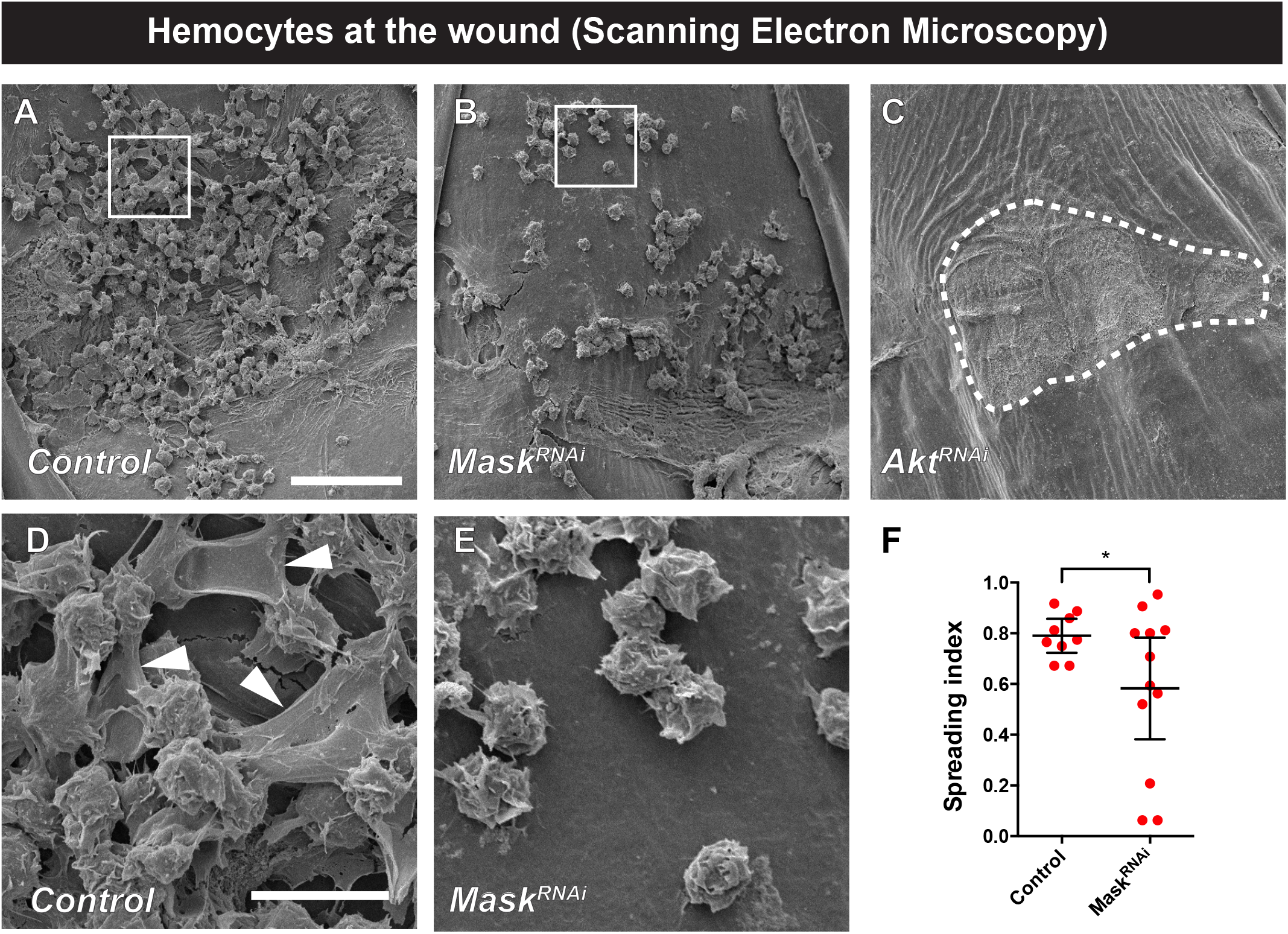
Mask is required for hemocyte spreading at wound sites. (A-E) Scanning electron micrographs of wounded and dissected third instar larvae of the indicated genotypes to visualize wound-adherent blood cells. (A) Control. (B) *UAS-Mask^RNAi^*. (C) *UAS-Akt^RNAi^*. (D) Control-closeup image of white box in (A). Arrowheads indicate hemocyte lamellae. (E) *UAS-Mask^RNAi^*-closeup image of white box in (B). In all panels (A-E) the bare larval cuticle and cell debris is underneath the attached hemocytes (see outlined region in (C) which lacks attached hemoctyes). (F) Quantitation of hemocyte spreading index at wound sites in control and *UAS-Mask^RNAi^*-expressing larvae. Unpaired *t*-test.

## Discussion

In this study we establish a new role for Pvf/Pvr signaling in regulating wound-induced blood cell spreading at the larval stage. Several lines of evidence suggest that the Pvf1 ligand and its Pvr receptor are required for blood cell spreading. First, blood cells in Pvf1 mutants show a rounded morphology at wound sites, unlike the typical spread morphology in controls. Second, Pvf1 can directly induce blood cell spreading *in vitro* in a manner that depends upon function of Pvr. Finally, Pvr hyperactivation promotes hemocyte spreading in primary cultures of larval hemocytes. Together these loss- and gain-of-function experiments strongly suggest that Pvf1 and Pvr are required for blood cell spreading. In this study we also developed a new screening platform to try to identify genes that might function downstream of Pvr in the various wound responses for which it is required. This genetic screen for suppression of Pvr-induced lethality identified a number of genes, some of which have strong phenotypes affecting WC, hemocyte spreading, or both. Below, we discuss the implications of these findings for wound-induced hemocyte responses, the diversity of Pvr signaling effects in different cell types, and the architecture of signaling downstream of Pvr in different wound-responsive cell types.

Larvae possess a population of circulating hemocytes that are distributed throughout the open body cavity and patrol for tissue damage (Brock et al. 2008). Hemoctyes that happen to bump into the wound adhere and spread (Babcock et al. 2008). Our work here suggests that adhesion and spreading are separable phenomena because in Pvf1 mutants and in larvae expressing *UAS-Mask^RNAi^* in hemocytes, attachment to wound sites occurs normally though subsequent spreading at the wound surface does not. In both of these genotypes the circulating hemocyte populations appear qualitatively normal. In the fly embryo, Pvr and several of its ligands are required for survival (Bruckner et al. 2004) and for developmentally-programmed hemocyte migrations (Parsons and Foley 2013; Wood et al. 2006) but not for recruitment to wounds (Wood et al. 2006). Other signaling pathways such as TNF are rquired for an invasive-like transmigration near the embryo head (Ratheesh et al. 2018). The differnatial role of Pvr and its ligands in embryos and larvae highlight another dimension to the interesting stage-specific differneces in hemocyte recruitment to damaged tissue (Brock et al. 2008; Ratheesh et al. 2015). There are other contexts besides wound-induced inflammation where hemocytes adhere to both normal and foreign cellular surfaces in *Drosophila*. These include sessile compartments (Bretscher et al. 2015), transformed tissue (Pastor-Pareja et al. 2008) and parasitic wasp eggs (Russo et al. 1996; Williams et al. 2005). It will be interesting to see in future studies if Pvf/Pvr signaling also plays a role in these events.

In addition to its roles in various developmental processes (Garlena et al. 2015; Harris et al. 2007; Ishimaru et al. 2004; McDonald et al. 2003), Pvf/Pvr signaling is required for a diverse array of tissue damage responses including epidermal WC (Wu et al. 2009), mechanical nociception (Lopez-Bellido et al. 2019) and larval hemocyte spreading during inflammation (this study). Both *in vitro* using S2 cells (Friedman and Perrimon 2006) and *in vivo* using glial cells (Kim et al. 2014) Pvr signaling screens have been carried out in other contexts. However, it has been a challenge to identify downstream Pvr signaling components that function in WC due to the lack of a pathway reporter that functions well *in vivo*. To circumvent this, we designed a genetic suppressor screen that exploits the fact that overexpression of Pvr in the larval epidermis is lethal (Wu et al. 2009). The reasons for this lethality are not clear but could potentially be related to a general hyperactivation of the epidermal WC response. If this hypothesis were correct, it might be expected that most identified lethality suppressors would also be required for WC. This was not observed. While a substantial set of the lethality suppressors were not found to affect WC, three factors - Ras, Mask, and MKK3 – did affect WC. This divergence between suppressors and WC genes could indicate a role for Pvr in maintaining the integrity or survival of the larval epidermis. Indeed, some of the genes found here overlap with Pvr signaling components found to be important for hemocyte survival (Sopko et al. 2015).

The three suppressors of Pvr-induced lethality that were also found here to be required for WC include Ras, a small GTPase; MASK, an adaptor protein required for RTK signaling in other contexts (Smith et al. 2002); and MKK3, a Map kinase kinase (Han et al. 1998). Epistasis analysis (overexpression of putative downstream Pvr genes in a Pvr-deficient background) revealed that only those components very close to Pvr in the presumed signaling cascade (Pvr itself and the Ras GTPase) were capable of partially rescuing the WC defect resulting from loss of Pvr. This could suggest that the Pvr signaling is performing multiple functions during WC and there is a split in the cascade downstream of the receptor (between Pvr/Ras and Mask/MKK3).

Interestingly, the Pvr suppressors found to be required for hemocyte spreading only partially overlap with those found required for WC. This is perhaps not too surprising since WC is a collective cell migration orchestrated by an epithelial tissue whereas hemocyte spreading is an individual change in morphology occurring in mesodermal cells. In summary, Akt is uniquely required for spreading *in vitro*; MKK3, Ras and Ck1α are only required for epidermal WC; and Mask is required for both *in vitro* and in vivo spreading and WC. These results suggest the signaling cascade downstream of Pvr differs in the two cell types and it will be interesting, now that genes are identified, to probe how these differences interact with the cytoskeletal architecture to achieve the observed changes in cell morphology.

## Acknowledgements

We thank Drs. Daniel Babcock and Amanda Brock for early work on wound-induced inflammation in our lab, Dr. Adriana Paulucci-Holthauzen and the MDA Department of Genetics microscopy facility for light microscopy assistance and Kenneth Dunner at the UT MD Anderson High Resolution Electron Microscopy Facility for electron microscopy assistance. Work in the Galko lab is supported through a “people not projects” mechanism-R35GM126929 of NIGMS. C.-R.T was supported by an American Heart Association predoctoral fellowship (16PRE30880004). AJ, NS, and JDC were supported by the CPRIT CURE Summer Undergraduate Research Training Program at MD Anderson Cancer Center (RP170067). Drs. Swathi Arur, George Eisenhoffer, and Galko laboratory members read and commented on the manuscript.

## Competing Interests

No competing interests declared.

---

**Figure S1.**
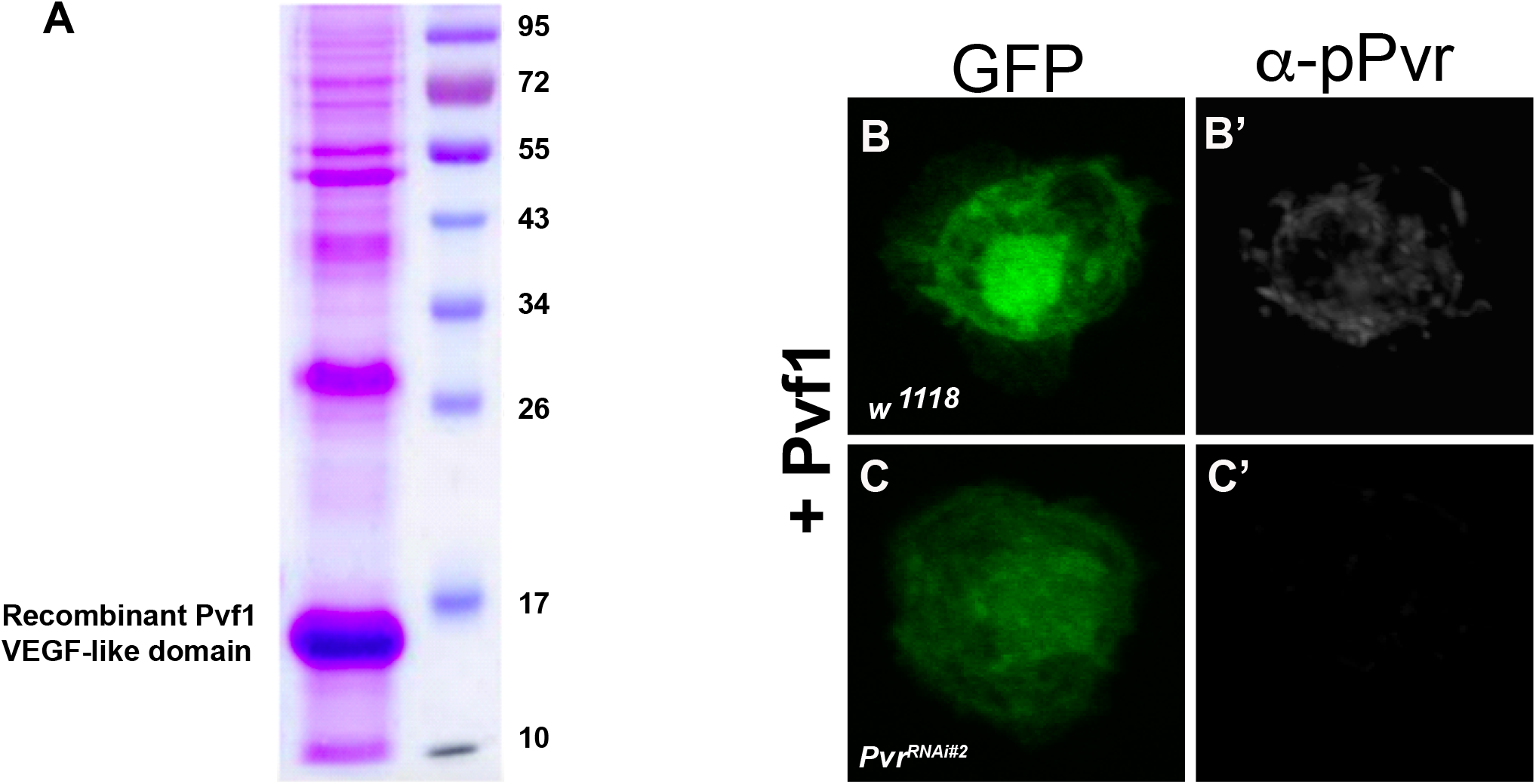
Verification of expression and activity of enriched Pvf1. (A) Enrichment of Pvf1 protein. Bacterial extract from cells overexpressing the Pvf1 VEGF-like domain (see methods). A large band at the expected MW of ~15 kD is observed. (B-C’). Hemocytes isolated from third instar larvae (genotype) +/- *UAS-Pvr^RNAi#2^* were plated for one hour, treated with enriched Pvf1 protein, and visualized with the *UAS-GFP* lineage marker (green) (B-C) or immunostained with anti-phospho-Pvr (white) (B’-C’). Only control hemocytes lacking the *UAS-Pvr^RNAi#2^* transgene show anti-phospho-Pvr staining upon addition of Pvf1.

## Supplemental Table 1. Flies used in this study

Please note the genotype of sex chromosome is simplified. The actual genotypes for the sex chromosome could be mixed, depending on the source RNAi collection, UAS transgenes, and larvae of both sexes were pooled and tested.

